# Computational modelling of toroidal membranes at division and fission sites

**DOI:** 10.1101/2025.05.02.651843

**Authors:** Chris L.B. Graham, Matyas Parrag, Phillip J. Stansfeld, Christopher D. A. Rodrigues

## Abstract

The formation of specialised membrane architectures is fundamental to biological processes. Computational modelling of membrane structures offers a means to unravel complex molecular mechanisms that remain inaccessible through *in vitro* or *in vivo* approaches. Among these architectures, toroidal membranes, otherwise known as fusion pores or fission pores, occur during the final stages of cell division, exocytosis, endocytosis, and membrane fission; including during the phagocytic-like engulfment of bacterial endospores. Here we have designed ‘MemTorMD’, a computational simulation pipeline based on the TS2CG methodology, to enable coarse-grained modelling of free toroidal membranes with biologically-relevant dimensions. This approach allowed us to determine the lipid properties required for stable toroidal membrane formation, providing a biologically accurate simulation surface. Furthermore, we demonstrate the applicability of this approach to investigate the function of proteins that localise at toroidal membranes, and drive molecular events such as membrane fission by a prokaryotic fission protein. Collectively, our work provides insight into membrane and protein behaviour at toroidal membranes and provides a platform for the development of more complex simulations involving additional molecular components at membrane toroids present in a range of processes across life.

## INTRODUCTION

Computational simulations of membrane structures can provide mechanistic insight into a wide array of biological processes, particularly those dependent on specific membrane architecture. However, most simulations of proteins within membranes – whether at coarse grained or all atom resolution – are performed using flat membranes rather than in curved, biologically relevant surfaces (1–5). Thus, to better capture the molecular mechanisms requiring curved membrane surfaces, advanced computational methods are needed to design and simulate these structures. Here, we develop and validate a versatile and reproducible computational method for free simulation of toroidal membranes through coarse grained molecular dynamics (CGMD) simulations, focusing on events that occur during cell division and bacterial endospore formation.

Toroidal membranes are central to various biological processes across prokaryotes, eukaryotes and archaea. For example, toroidal membranes are formed during endocytosis and exocytosis, where toroidal membranes eventually undergo fusion and fission events that allow the uptake, or release, of membrane vesicles into and from cells, respectively (6–9). Longer toroidal membranes, flanked by extracellular walls, are formed during intercellular communication, such as those found in plasmodesmata of plant cells (10). Importantly, toroidal membranes are critical to the process of cytokinesis, creating the ‘fusion pore’, where invaginating membranes slowly come together during the final stages of cell division and eventually result in cell separation(11,12).

Previous studies have advanced the simulation and characterization of biologically relevant toroidal membranes . However, as with many computational simulations requiring curved membranes, toroidal membranes have generally been generated using dummy particles, or from fragmented flat membranes(6,8,13–15). Dummy particles reinforce the shape of the membrane during simulation independent of energy in the system, thus constraining lipids to a high energy state(16). For example, recently the toroidal membranes associated with the folds of the inner membrane of mitochondria have been simulated to understand lipid dynamics and localisation on these curved surfaces supported by dummy particles (13,14). Whilst we have learnt a great deal using these setups, this approach intrinsically limits the range of biological questions that can be asked of toroidal membranes, especially if we consider that toroidal membranes come in different shapes and sizes that alter their stability, curvature and lipid density, and therefore could also affect protein behaviour, depending on the organism, the organelle and fundamentally the biological process (10,17–20). Importantly, the removal of dummy particles results in an unrestrained system and causes shape deviations from an initial configuration towards lower energy states, which in some case involves collapse of the membrane architecture (16).

Recently the TS2CG package has been used to orient coarse grained phospholipids in vesicles using user-defined parameters with and without dummy particles (21). Specifically, a 3D-triangulated surface is assigned with lipid molecules through pointillism transformations, generating for example, a three-dimensional model of a membrane vesicle (21). Our closer inspection of this work highlighted that toroidal membranes connecting vesicles can be simulated using this approach.

Here, using a modified version of the TS2CG package which enables free membrane edges(22), along with custom scripts and parameters reproducible within *Blender .*stl design software, typically used in game design and animations (23), we simulate toroidal membranes of varying dimensions and lipid densities without requiring dummy particles. With a focus on bacterial toroidal membranes, we develop a computational method for generating toroidal membranes with diverse dimensions, lipid densities and composition. We use nano-to-microsecond scale coarse grained simulations to demonstrate the effects of lipid membrane composition and lipid concentration on the stability of the toroidal membrane. We then apply this framework to test the behaviour of proteins known to exist at toroid sites. Specifically, we investigate the role of a highly-conserved prokaryotic membrane fission protein, FisB, which is localized at a toroidal membrane in sporulating bacterial cells and is required for membrane fission (7,24). Collectively, our method provides a platform to study lipid and protein behaviour at toroidal membranes. It also allows for the development of more complex simulations in the future, including additional cell envelope components, and facilitates the investigation of many biological processes that occur at toroidal membranes, including division.

## METHODS

### MemTorMD pipeline: Coarse grained lipid toroid simulation

MemTorMD is a python script executed in Google Colab, which relies on the ‘Triangulated Surface to Coarse Grained’ (TS2CG) script. The TS2CG software we use can calculate boundary exceptions; in other words, it can be used on small systems that are not complete such as two parallel membranes, as opposed to vesicles. All scripts rely on a .tsi file to function. The pyc2tsi Blender script was used to create three .tsi file a flat membrane. The .tsi files are transformed by scaling the x, y and z dimensions of the vertices to alter toroidal membrane height and pore diameter. The default simulation used neutralising salt conditions with NA ions, dependent on POPG number to counteract the negative charge of the POPG. Finally the simulation after equilibration is further stabilised for 2000 frames dt = 0.0001 picoseconds to reduce clash issues, before finally being simulated. The default simulation conditions, and scripts, are attached in Supplementary Information and available on the MemTorMD pipeline. A user uploads a protein or up to two pdb files of choice, then determines the location of insertion of each protein, the lipid composition, salt concentration, the degree of x, y and z transformation therefore toroid size are user determined.

### Co-evolution quaternary interaction resolution

Co-evolutionary contacts within FisB were calculated using the GREMLIN algorithm (25). MMSEQ2 was used to create the sequence alignment with scripts available here Co-evolution suite (26). The AlphaFold2 structure of monomeric FisB, and 20-mer FisB oligomer, were then compared to potential multimeric AlphaFold3 structures by their respective contact maps, with a carbon backbone threshold set to 12 Å irrespective of chain number, to show a multimeric contact map interface. The co-evolution pairs which agreed with the monomer were labelled in blue, and in multimer only in red, to reveal multimer only interaction points.

### APL calculations in 3D lipid environments

Calculation of area per lipid (APL) involved two main steps: calculation of surface normals at each lipid and Voronoi tessellation of membrane patches (27). We can obtain a patch of lipids centred on a given lipid L, by assessing at all lipids within 30 Å. In this case the position of a lipid is the XYZ-coordinates of a user specified atom contained within the lipid. SL denotes this set of lipids (**S**et of **L**ipids). SL will sample a curved surface, which is inherently 3D. 2D Voronoi tessellation was used for simplicity and computational efficiency, hence a 2D set. TL (**T**esselated **L**ipids), is constructed from SL by projecting each bead in SL. Voronoi tessellations is performed on TL and the area corresponding to L is assigned as the area of L. As can be seen, calculation of the normals is vital for accurate APL calculation. The lipids sample a surface, however accurate estimation of this surface from lipid positions is difficult and expensive due to the sparsity and noise of samples. The normal for a lipid L can be estimated efficiently as the average direction of nearby beads to the central bead representing L. The script ‘Area_per_lipid_calculator.py’ available on the GitHub repository, was used to implement this calculation on supplied .gro files of the tested segments and/or separated leaflets.

### Multimer prediction by AlphaFold3 and AlphaFold2

The 1-184 SpoIIIE sequence was modelled as a multimer by uploading its amino acid sequences to the AlphaFold3 servers (Fig. S9A). The FisB 20-mer used in the simulations was predicted using the 16-216 FisB sequence by AlphaFold2(Fig. 5C) and compared to AlphaFold3 models with alternative stoichometry of the same input (Fig. S6A).

### Protein insertion by lipid replacement

Proteins were inserted into toroidal membranes manually. For FisB, a toroidal membrane with a 14 nm pore, with set lipid composition (2% CDL1/ 20% POPG/ 78% POPE) was set up using MemTorMD without proteins and simulated for 20 ns. The waters and ions were then removed, and a coarse grained FisB 20-mer positioned into the pore after 20 ns. All atoms within 1.5 Å of the 20-mer were then removed for lipid-replacement using PyMOL. Finally, a script calculated the number of lipids in the file, and the FisB Martini files added to the topol.top file, for simulation by the standard MemTorMD script.

### Tension calculations

To calculate tension, we made a simple assumption in the context of setting-up the simulation conditions, relative to input or output APL. Input APL is the APL introduced into the TS2CG framework and output APL is the APL calculated from the Voronoi tessellations after 10 ns during simulation. In this context, a zero-tension output APL arises when the APL remains constant across a range of input APL values which then goes on to increase output APL linearly above this range, as shown in Figure S1 (green region). Lipid tension is calculated from the change in percent APL over this optimal zero tension surface area (optimal APL) and an approximate 0.28N change in percent APL (τιAPL) (7). We assume an area compressibility modulus of K A = 0.28 N m^−^1.

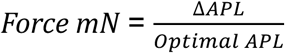

## RESULTS

### Design and scaling of reproducible toroidal membrane structures based on bacterial membranes

Although toroidal membranes have been simulated through a variety of methods, these existing methods did not always create a free-standing toroidal structure directly (6,8,13–15). Instead, membrane surfaces were often manipulated to induce a toroidal structure by a ‘dummy’ bead barrier that forces a toroidal membrane into shape (14). With the recent increase in computational power that allows for coarse grained simulations over microsecond time periods, and new tools that allow insertion of lipids into user defined 3D structures without the need for ‘dummy’ bead barriers, we set out to develop a re-usable pipeline and computational framework to study toroidal membrane, principally in bacterial membranes.

To build a toroidal membrane, we devised a simple framework (Fig.1) based on previous work utilizing the triangulated surface to coarse grained simulation methodology (TS2CG) (21). First, we designed the toroid shape through use the *Blender* (23) performing triangulation, screw and mirror transformations on a curve drawn on a 2D surface to create the triangulated 3D surface of a toroid (Fig.1Ai-iv). The triangulated surface was then stretched using multiplication factors in the X, Y and Z planes to create a variety of toroid dimensions. The resulting architecture, after pointillism was modified to produce two surfaces representing the two membrane leaflets of a toroidal membrane (Fig. 1B). We then utilised the coarse grain lipid insertion of a modified TS2CG programme (22) to insert lipids onto this surface at a user defined Area per Lipid (APL) (Fig.1C). Finally, the resulting set-up was then sliced by MDAnalysis (28) to remove excess atoms and to create a cube cell for simulation by GROMACS (29).

**Figure 1:**
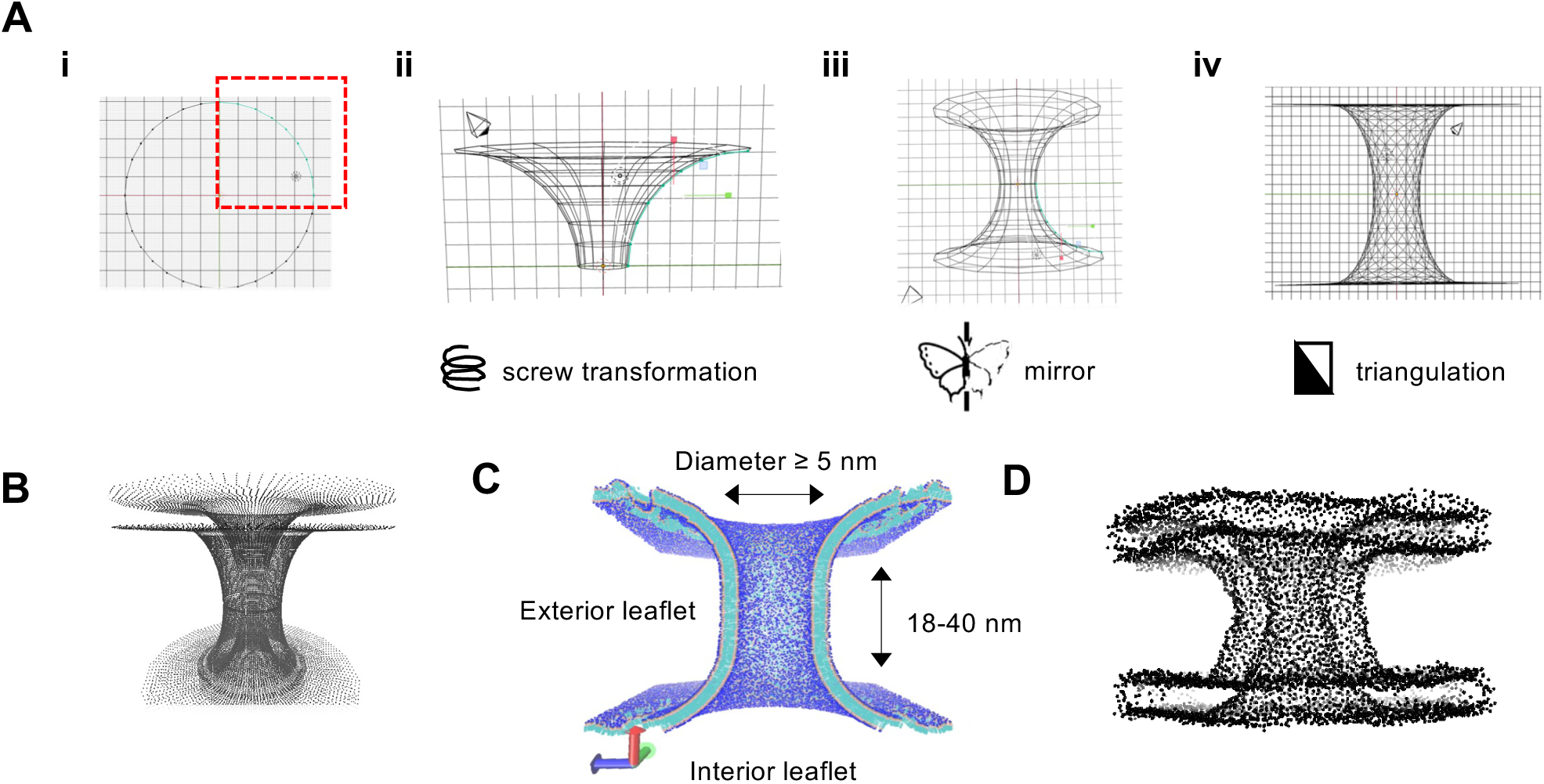
Computational approach to develop a toroidal membrane. **(A)** Steps utilized for toroid stl preparation within Blender: (i) the corner of a circle is created on the xy plane; (ii) vertexes of circle corner are transformed to dimensions 4.5 nm in xyz and future pore inside diameter is determined by screw distance to y axis, using the blender screw object modifier; (iii) the resulting vortex is converted to a toroid through blender mirror modifier; (iv) final .stl preparation and export through use of ‘alternative triangulation’. **(B)** Visualization of points prepared by Pointilism: the TS2CG stl2tsi .pyc script converts the .stl to a tsi file. PL pointillism 0.4 bond length, 4 nm leaflet produced from .tsi coordinates with a resize of 2, 2, 2. **(C)** Visualization of initial TG lipid population performed with custom .str file creating a .gro file assigned with custom lipids along .tsi coordinates. **(D)** Phosphate headgroups (black) of a 20 nm high and 4 nm diameter toroidal membrane pore after 10 ns. Cylindrical lipid placements are converted to box for parallel simulation in MD analysis by deleting outlying atoms. Custom .topol files are created by searching for lipid populations in that new box; Simulations were set to 310K, 1 bar, and then simulated.

To adapt the above framework to bacterial cells, we considered the dimensions of a toroidal membrane based on what has been reported for the model organisms *Bacillus subtilis* and *Escherichia coli* (17–19). In *B. subtilis and E. coli,* cryo-EM data suggest that the intermembrane spaces that separates the two leaflets of the toroidal membrane during cell division varies and can be between 20-40 nm in height, depending on the stage of life cycle(18,19). These dimensions include the height of two lipid bilayers and the space that separates them, which in the biological setting contains peptidoglycan (30). Since DNA is often present at the toroidal membrane during the final stages of cell division, we also considered this constraint in defining the diameter of the toroidal membrane pore, and set this size to 5 nm, which is large enough to accommodate two stands of double-stranded DNA. Initially, we adapted the composition of the membrane to reflect a simple system containing 100% POPG, but then later adjusted lipid concentrations to those typically observed *in vivo* in *E. coli* (2% Cardiolipin, 78% POPE, 20% POPG) (31). Based on these biological parameters, we were able to create a coarse-grained model of a toroidal membrane during the final stages of division, initially for 10 ns (Fig. 1D).

To improve usability and provide a user-friendly interface to the community, the resulting TS2CG pipeline has been made accessible on Google Colab and GitHub, with the necessary scripts for simulation of *E. coli* and *B. subtilis* toroidal membranes, including Martini insertion of proteins and inout of lipid concentration – we called this interface “MemTorMD”. Furthermore, the MemTorMD pipeline is customisable to accommodate simulations involving toroidal membranes of different size and also allows for simulation of vesicles. For the remainder of this work, we have applied the MemTorMD pipeline to understand the dynamics of a toroidal membrane in coarse grain and compare this with existing *in vivo* data, including the observation of simulated proteins.

### The stability and size of toroidal membranes in-silica is dependent on lipid density

In our simulations, although the toroidal membrane changed morphology within the first few nanoseconds, it then settled into a defined toroidal morphology (Fig. 2B). Toroidal membranes were considered stable during simulation when lipid number became constant within a fixed area in the cylindrical portion of the toroidal membrane (Fig. 2B & C). Under this measure of stability, we determined that the toroidal membrane stabilized at approximately 10 ns into the simulation (Fig. 2B & C).

**Figure 2.**
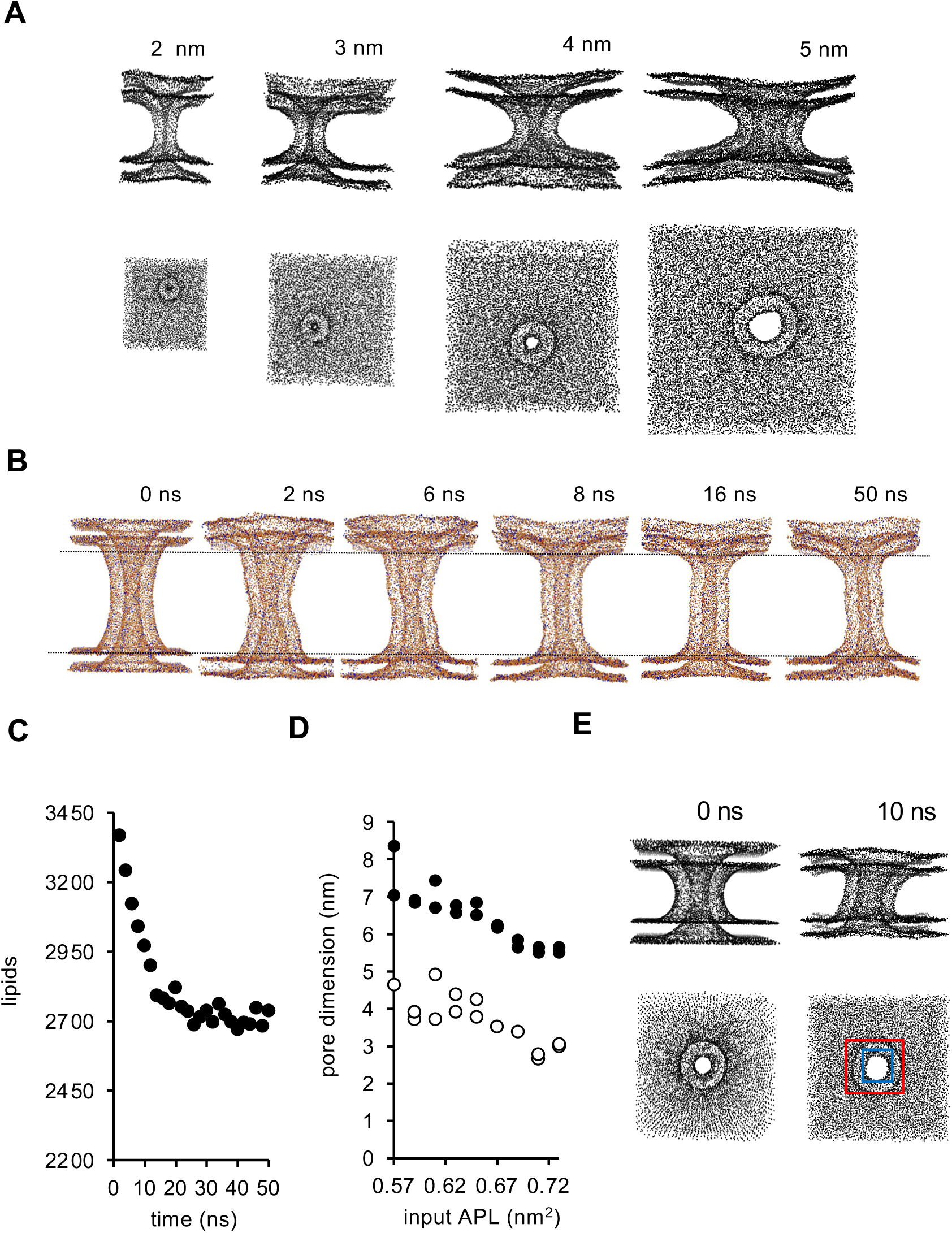
Pore size, toroidal membrane stabilization and initial simulation observations. **(A)** Phosphate headgroup visualization of a range of toroidal membranes, simulated at a stable APL (0.65 nm^2^ input APL) after a 500 ns timeframe. Internal pore diameter indicated on the top right (**B)** Toroidal membrane morphology during initial 50 ns stabilization, in a 40 nm high toroidal membrane with POPE:POPG:CDL1 lipids; simulation conditions were 1 bar, 310K, 5e^−6^ Box, compressibility and 0.70 nm^2^ input APL. **(C)** Initial lipid equilibration during simulation of toroidal membrane, determined at defined location within the cylindrical portion of the toroidal membrane (region between the black lines). (**D)** Relationship between input APL differences and toroidal membrane pore dimensions: interior leaflet diameter (white circles), exterior leaflet diameter (black circles); POPG only membrane. (**E**) Exemplification of toroidal membrane pore diameter measurement after 10 ns stabilization; interior leaflet (blue box), exterior leaflet (red box).

However, to determine an APL that would result in a stable toroidal membrane for POPG only membranes, we trialled nanoscale simulations with a toroidal membrane of a fixed dimension (40 nm height), at a range of input APLs (Fig. S1). One of the observations we made was that input APL using TS2CG did not reflect the actual output APL observed during simulation (i.e. output APL). To accurately measure APL during simulation, we designed a programme that could assign an output APL to toroidal membranes within a range of input APL (Fig. S1). Furthermore, with this programme we investigated the relationship between input and output APL and how it influences lipid tension during simulation (Fig. S1).

We found that the average output APL after 10 ns (i.e. after stabilisation) varied between 65.4 Å^2^ and 71.3 Å^2^ (Fig. S1A) and we made four major observations:1) at low input APL (<55 Å^2^), an increasing range of output APLs were observed and gave rise to new curvatures within and around the cylindrical portion of the toroidal membrane (Fig. S1A; *grey region*); 2) when the input APL was between 55 Å^2^ and 63 Å^2^, the output APL remained relatively constant suggesting an equilibrium and a near zero tension lipid environment (Fig. S1A – *green region*); 3) as input APL increased beyond 63 Å^2^ and up to 73 Å^2^, we observed a near linear increase in the output APL and that the toroidal membranes generally appeared more stable (Fig. S1A – *blue region*); and 4) above an input APL of 73 Å^2^, the toroidal membrane became unstable and teared apart at the edges (Fig. S1A – *red region*). Based on this, we inferred that input APL values between 63 Å^2^ and 73 Å^2^, cause an increase in membrane tension and changes in toroidal membrane shape (as seen in Fig. S1A – *blue region*). Interestingly even after 200 ns, in simulations where there was no tearing apart at the simulation edges (73<Å^2^), the pore of the toroidal membrane remained open (Fig. S1 & 2), suggesting that the membrane is relatively stable on its own and does not collapse or fuse spontaneously. Furthermore, calculating tension, by applying the formula described in the Material & Methods, suggests that toroidal membrane stability is achieved between 0 and ∼8 mN/m. Thus, by comparing input APL and output APL, we were able to identify conditions where toroidal membrane remain stable during simulations without the need for force beads.

Next, to test if MemTorMD can be used to study toroidal membranes of other dimensions, we investigated the relationship between input and output APL on a toroidal membrane set to 20 nm height, 4 nm pore diameter and 100% POPG (Fig. S2). Here we observed a less linear relationship between input and output APL but nonetheless identified a range of input APLs that results in stable toroidal membranes (Fig. S2). Interestingly, while conducting these simulations we found that toroidal membrane dimension partly depends on the input APL (Fig. 2D). Indeed, a low input APL (<0.62) led to increased toroidal membrane pore diameter and instability, leading to deformed structures (Fig. S2B). In contrast, higher input APLs (>0.73) decreased toroidal membrane pore diameter (Fig. 2 & S2B). Collectively, based on these observations examining the relationship between input and output APL (and tension), were could confidently define a range of APLs that generate stable toroidal membranes.

### Simulation of a range of stable toroidal membranes with different pore sizes

Based on the above, and demonstrating the versatility of the MemTorMD script, we then created of a set of stable toroidal membranes with pores of different diameters, between 1 nm and 20 nm wide with the 78%:20%:2% POPE:POPG:CDL1 configuration, by triangulated surface stretching of the 20 nm high toroidal membrane shown in Fig. 2A. Each toroidal membrane had different starting APL (between 0.62 nm^2^ and 0.71 nm^2^) which changed its overall shape. Interestingly however, most toroidal membranes with pore diameters between 1-20 nm could be simulated at 0.65 nm^2^ APL (Fig. 2A). Importantly, we did not observe spontaneous fission of the membrane in any of the stable toroidal membranes assayed so far (each with simulation lengths of 500 ns).

### Modelling bacterial toroidal membranes of biologically-relevant proportions

Next, we set-out to model and simulate three toroidal membranes that occur at different stages of *B. subtilis* life cycle: 1) a toroidal membrane delineated by thick peptidoglycan found during vegetative growth (medially positioned division septum) or sporulation (polarly positioned division septum), with an estimated pore height of 40 nm, based on cryo-ET data (17,19) (Fig 3Ai); 2) a toroidal membrane delineated by thin peptidoglycan found at the curved polar septum at the onset of septal peptidoglycan hydrolysis during a process called engulfment which internalizes the developing spore into the mother - estimated height of toroidal membrane is 20 nm, based on cryo-ET data (19)(Fig 3Aii); and 3) a toroidal membrane that occurs at the membrane fission event which completes engulfment of the spore by the mother cell (Fig. 3Aiii). Due to the absence of cryo-ET data showing the engulfment fission event, the toroidal membrane associated with fission was modelled to a 15 nm diameter pore and 20 nm in height (Fig. 3Aiii). Simulations over 5 µs showed that each type of toroidal membrane was stable. Stability was achieved at 0.67 nm^2^ APL for the 20 nm high toroidal membrane associated with polar division after peptidoglycan thinning (Fig. 3Bii), 0.71 nm^2^ APL for the 40 nm toroidal membrane associated with polar division (Fig. 3Bi) and 0.67 nm^2^ to 0.60 nm^2^ APL (APL dependent on the input toroidal pore diameter) for the larger toroidal membrane associated with engulfment membrane fission (Fig. 3Biii). These data suggest that toroidal membranes of dimensions similar to those encountered *in vivo* have potential for modelling *in silico* without dummy particles.

**Figure 3.**
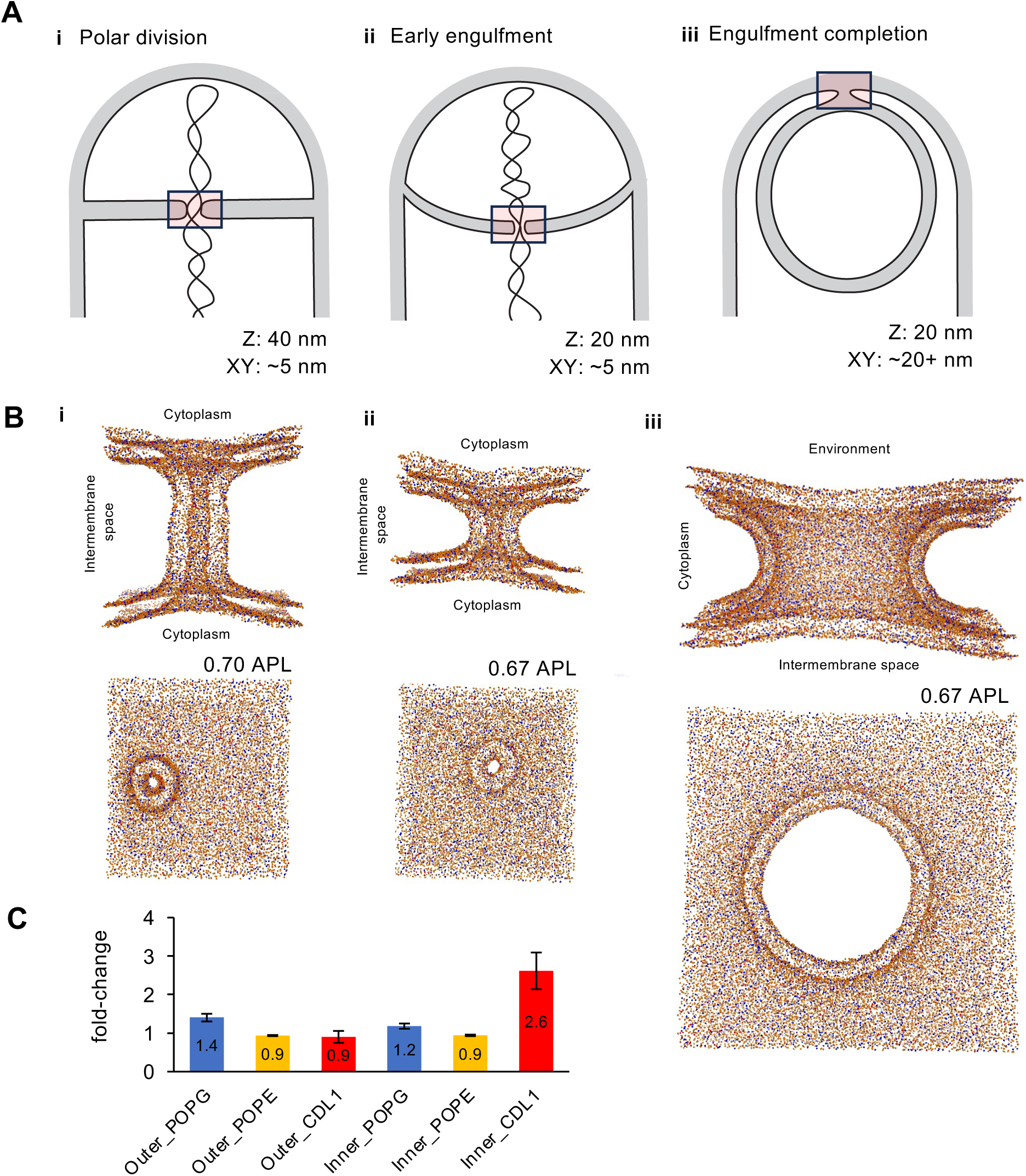
Lipid distribution, optimal APL and morphology of toroidal membranes during longer simulations. **(A)** Schematic illustration of toroidal membranes (inset red box) found during the *Bacillus subtilis* life cycle: during **(i)** polar division (thick septum), **(ii)** onset of engulfment (thinner, remodeled septum) and **(iii)** final stages of engulfment; dimensions are based on existing data as described in the text. **(B)** Phosphate headgroups visualization in the life cycle stages described in (A): **(i)** 40nm, 40.5×40.5 nm box**, (ii)** 20 nm, 40.5×40.5 nm box and **(iii)** 70.5×70.5 nm box. Simulation conditions were 80% POPE/ 18% POPG/ 2% CDL1 ,1 bar, 310K, 5e^−6^ Box, compressibility. Red-CDL1, Blue-POPG, Orange-POPE. (**C)** Lipid migration to the interior and outer leaflets of toroidal membrane in a 20 nm high, 4 nm diameter toroidal membrane after 10 *μμ*s; 310K, 1Bar and starting lipid concentration 20% POPG, 78% POPE, 2% CDL1 (n=3).

### Toroidal membranes pores have subtle and reproducible local differences in lipid composition

*In vivo* there are a variety of lipid types with chemically distinct headgroups and tails(31–34). This facilitates lipid domain formation over time (32,35). Given the stability of our toroidal membranes during simulation, we wanted to know if we could observe the formation of lipid domains during simulation, or at the very least observe differences in local lipid abundance within the toroidal membranes. For example, cardiolipin has been shown to localise at areas of high curvature present during cell division in *E. coli* (36), and in curved membranes in simulations (15,36,37). Thus, based on the curvature and extreme APL values found within the toroidal membranes we have simulated thus far, it seemed likely that cardiolipin would become enriched at certain locations within the toroidal membranes.

We examined the toroidal membrane for changes in lipid composition, both in the interior and exterior leaflet, after a 10 µs simulation of an *E. coli* membrane (<u>Video 1</u>). Our simulations showed that by 1 µs, full mixing of the lipids had occurred (Fig. S3B). Lipid domains were observed at 10 µs; however, no noticeable domains formed within the pore region of the toroidal membrane (Fig. S3A). Nonetheless, in a toroidal membrane with a height of 20 nm and 0.67 nm^2^ APL, we observed significant changes in cardiolipin concentration over the course of the simulation (Fig. 3C). Cardiolipin concentration significantly increased (over 2-fold; from 2% to 5%) in the inner leaflet of the toroidal membrane over the course of the simulation (Fig. 3C). Conversely, on the exterior leaflet there was no significant change in cardiolipin concentration. The observation that cardiolipin accumulates in interior leaflet and not the exterior leaflet, is likely due to differences in curvature between the two leaflets. Thus, consistent with biological and *in silica* data showing that cardiolipin is increased in bacterial membranes of high membrane curvature, these results suggest that computationally simulated toroidal membranes by MemTorMD can reproduce existing observations relating to lipid localization and domain formation.

### Toroidal membranes pores have local differences in lipid density

The above datasets suggest that APL in a toroidal membrane is different to that of a flat membrane surface. Previous work suggests that in artificially curved systems, APL is dependent on lipid surface curvature (37). However, in our own system, where the toroidal membrane is stable due to variations in tension, it remains unclear how the APL changes across toroidal membranes. To answer this question, we simulated toroidal membranes containing POPE:POPG:CDL1 and with pore diameter between 1 nm and 33 nm, with fixed height and APL (Fig. 4). At the end of each simulation, we accessed APL along the Z-axis of the interior and exterior leaflet of the toroidal membrane (Fig. 4A, Fig S3). We found that APL changes on both leaflets and is dependent on pore diameter. Interestingly, APL on the interior leaflet of the toroidal membrane pore region was lower than planar membrane APL (Fig. 4A(2,38), with values between 0.48 nm^2^ and 0.55 nm^2^. In contrast, in the exterior leaflet, the APL varied between 0.63 nm^2^ to 0.90 APL nm^2^ (Fig. 4). We also found that the extremity of these values was dependent on the morphology of the toroidal membrane, with thinner and longer toroidal membranes having more divergent APL than shorter and wider ones (Fig 4C). This may reflect reduced tension and curvature across larger surfaces, as suggested previously (16).

**Figure 4.**
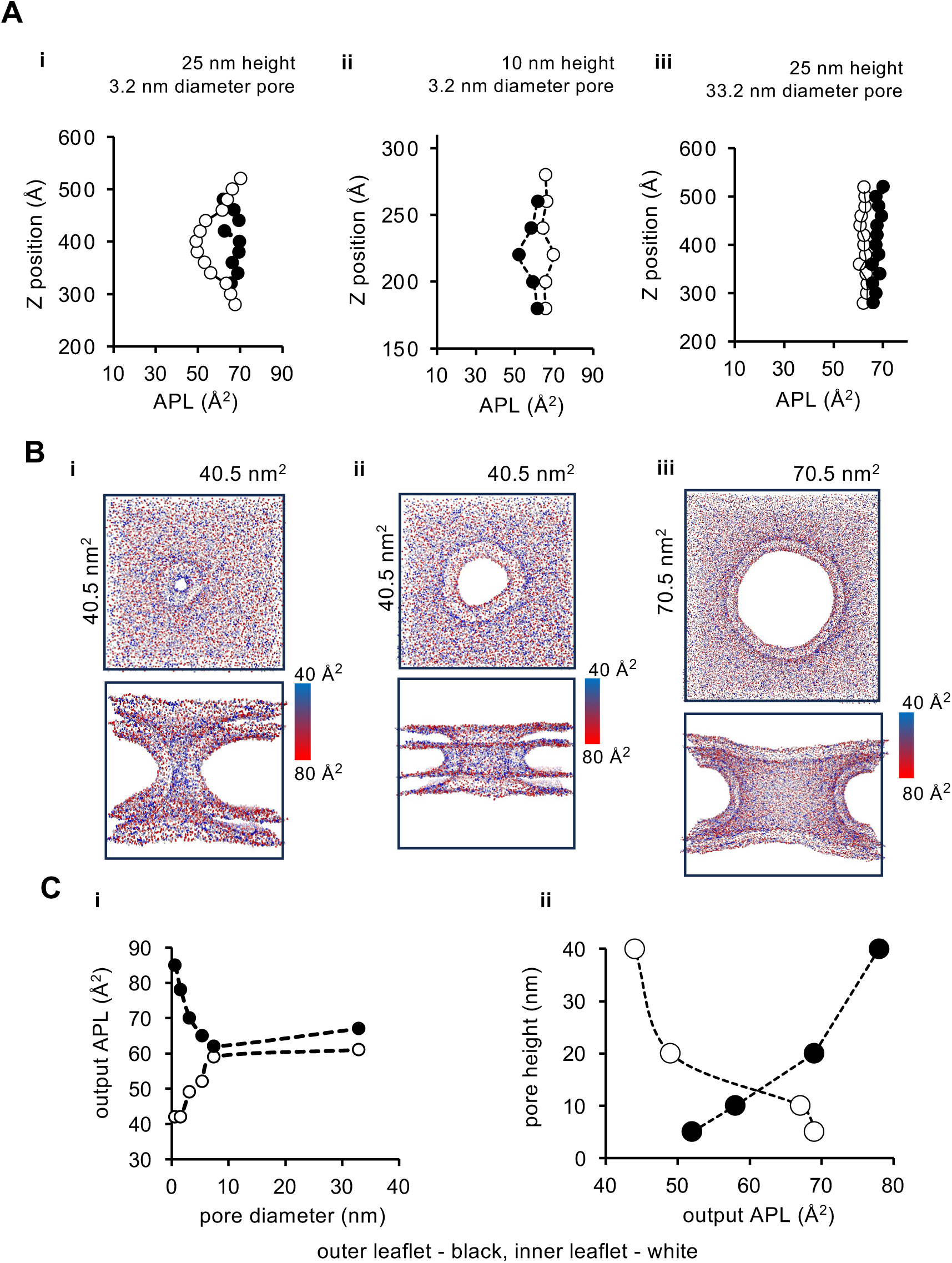
Lipid domains, APL and headgroup concentration differences within the pore region of the toroidal membrane. Simulation conditions were 1 bar, 310K, 5e^−6^ Box, compressibility. Red-CDL1, Blue-POPG, Orange-POPE **(A)** Area per lipid along the Z-axis of toroidal membranes with different dimensions, determined along 20 Å interval Z slices. (**i)** 3.2 nm pore, 25 nm height **(ii)** 3.2 nm pore, 10 nm height**, (iii)** 33 nm pore, 25 nm height; white circles - interior leaflet, black circles - exterior leaflet. (**B)** APL density by phosphate position after 500 ns. Scale is Blue – 40 nm APL to Red 80 nm APL. (**C)** Area per lipid depends on pore diameter **(i)** and height **(ii);** APL was measured in the interior leaflet (white circles) and exterior leaflet (black circles).

**Figure 5.**
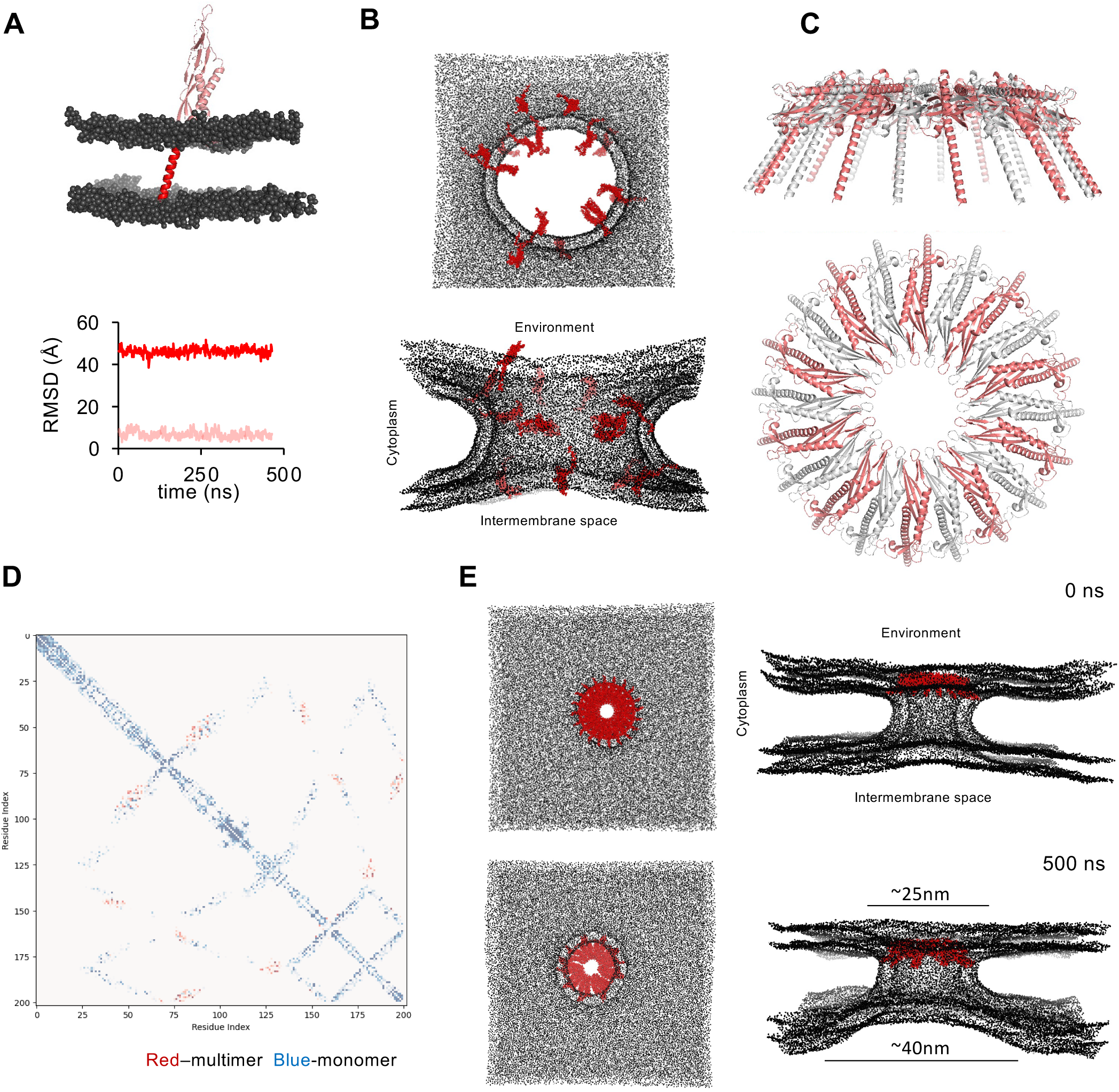
FisB protein complex formation within a toroidal membrane. **(A)** Monomeric FisB and flexibility of its transmembrane domain (red), relative to the extracellular globular domain (pink). Simulation for 500 ns. Graph below shows the root mean square deviaton (RMSD) for the transmembrane (red) and globular domain (pink). **(B)** Clustering of FisB at the toroidal membrane pore; 0.62 nm2 APL prepared by MemTorMD, 16 FisB molecules. Simulation for 500 ns. **(C)** AlphaFold rendering of FisB-20 mer; protomers are shown in red and grey and the Nterminal transmembrane was removed; Details of coevolution pairs present in the multimer are described in Fig. S6. **(D)** Overlay between co-evolution and predicted C-C backbone multimer/monomer contact maps suggests FisB-FisB specific interactions within a multimer contact. A 10 Å cutoff was used. **(E)** Coarse grained simulation of FisB-20 mer within a 10 ns pre-stabilised toroidal membrane, demonstrates that FisB clasps the toroidal membrane. 14 nm wide pore, 20 nm height, POPE 78%, POPG 20%, CDL1 2%, 310K, 1 bar, 0.5 *μμ*s simulation, diameter of pore at flat surface of each side indicated. Refer to Fig. S9 for additional tensions and timepoints.

We also assessed the effect of the changing the height of the toroidal membrane on APL, by stretching the .tsi file in the Z-dimension. Plotting APL against toroidal membrane pore height showed that APL in shorter pores was higher in the interior leaflet than the exterior leaflet, with the interior leaflet APL being comparable to that observed in flat membranes (Fig. 4Cii). However, APL on the exterior leaflet of toroidal membranes with shorter pores was lower, likely due to the extreme curvature of the exterior leaflet surface found on toroidal membranes with shorter pores (Fig. 4Bii). In contrast, the opposite was observed with toroidal membranes containing taller pores (Fig. 4Biii, Fig. S4E). Collectively, these data suggest that APL, and therefore tension, is a critical factor in determining the shape of a toroidal membrane: a highly concave architecture in a toroidal membrane leaflet causes a high APL, and highly convex architecture therefore causes low APL.

### Simulation of proteins during engulfment membrane fission

The above data characterize the basics lipid properties of computationally simulated toroidal membranes using MemTorMD. Next, we sought to examine if MemTorMD would allow examination of protein behaviour within toroidal membranes, as well as how proteins that exist at toroidal membranes affect the toroidal membrane itself. Of the bacterial proteins known to exist at the toroidal membrane during cell division, many are related to the regulation of peptidoglycan synthesis (39–43). Since our computational framework is currently limited to membranes, we sought to examine the behaviour of a protein that exists at a toroidal membrane and acts directly on lipids. We thus focused on FisB, a protein that is required for membrane fission during the final stages of spore engulfment in *B. subtilis* (Fig. 3Aiii and Fig.5) (7). Current data suggests that FisB accumulates at the site of membrane fission as a single focus, where it forms oligomers that bind to lipids, slow down lipid movement and create membrane tension at the toroidal membrane that lead to membrane fission(7). Thus, we considered FisB the ideal candidate to observe the dynamics of proteins and lipids at a toroidal membrane.

Since FisB is known to oligomerize *in-vivo* and *in-vitro*, we first tested if we could observe FisB oligomerization at the toroidal membrane during simulation. Previous work indicates that approximately 40 FisB molecules exist at the membrane fission pore and are required for efficient membrane fission; however, fewer FisB molecules are sufficient to mediate fission (as low as 6) but at a slower rate(7,24). We introduced 16 FisB monomers at random locations surrounding the neck of the pore of a toroidal membrane that was 20 nm in diameter and 20 nm high. Since the addition of proteins to the toroidal membrane framework alters the APL, there was expansion of the pore caused by torsion at 0.67APL. After 100 ns of the simulation, we observed that FisB began to form dimers (Fig. 5B, <u>Video 2</u>) and after 500 ns distinct FisB multimers appeared (Fig. 5B). Thus, FisB can oligomerize *in silico*, in the context of the toroidal membranes.

Although FisB has been shown to form oligomers of varying molecular weight *in-vitro* (7,24), the exact structure of a FisB oligomer at the membrane fission pore has yet to be established. However, quantitative analysis of fluorescence intensity of FisB-GFP fusions, suggest there are roughly 47± 20 FisB molecules at the fission pore. Interestingly, AlphaFold2 (44) predicts a ring-shaped FisB 20-mer (Fig. 5C), among other possible ring-shaped multimers. Closer inspection of this hypothetical FisB 20-mer suggests that these multimers are likely biologically relevant since residues G175, I176, I177 and I194 implicated in FisB oligomerization *in-vitro* and in FisB-GFP focus formation *in-vivo* (24), are located at, or near, the interface of FisB protomers within the 20-mer ring model (Fig S6C). Furthermore, and providing evolutionary evidence that the FisB 20-mer ring model is likely biologically significant, co-evolutionary analysis of residues (25) not identified as contacts required for the folding of the FisB, are present exclusively in these ring-mers including the FisB 20-mer. (Fig. 5D). Furthermore, this analysis suggests that the FisB 20-mer is better supported by quaternary co-evolution data, compared to other FisB multimer models (Fig 5D). Thus, whilst it remains unclear if a ring-shaped 20-mer is the *in vivo* structure of a FisB complex at the fission pore, existing *in vitro* and *in-vivo* data, and co-evolutionary analysis (Fig. S6B), support the idea that a FisB 20-mer could be involved in membrane fission event. Based on this, we sought to examine the behaviour of the FisB 20-mer in a toroidal membrane pore.

When considering how the FisB 20-mer ring could assemble into the toroidal membrane, we predicted that FisB’s head domain, implicated in oligomerisation and lipid binding *in vitro* (7), would exhibit some flexibility relative to the N-terminal membrane-spanning helix. Indeed, atomistic computational simulation in planar lipids confirmed the presence of a flexible hinge between the membrane-spanning helix and the head domain *in silico* and demonstrated a wide range of movement indicated by divergent RMSD values ∼10 v ∼45 over 500 ns (Fig. 5A, S6A). Thus, the flexible nature of the FisB head domain would allow FisB to oligomerise and adopt a conformation that aligns with the lipids in the pore of the toroidal membrane (Fig. 5A). Based on these considerations, we then inserted the hypothetical FisB ring shaped 20-mer into toroidal membranes represented by a MARTINI 3 coarse grained structure and examined the behaviour of FisB and the membranes during nanoscale simulations.

Interestingly, initial simulations of the FisB 20-mer in the toroid using MemTorMD led to hemi-fission-like events (Fig. S8i, Videos <u>5/6/7/8</u>). We considered the possibility that the multimeric FisB 20-mer configuration within the toroidal membrane likely interrupted the APL stabilisation steps at the start of the simulation, caused membrane instability and led to these hemi-fission-like events. To test this idea, we repeated the simulation with the FisB 20-mer, but allowed membrane equilibration for 10 ns without FisB, to reduce instability of the starting configuration. Then, we added FisB by lipid-replacement. In this scenario where the toroidal membrane was stabilised for 10 ns before addition of FisB, FisB initially maintained the toroidal membrane pore diameter in a stable manner (Fig. 5D, <u>Video 3</u>), but then began to act like a clasp, changing the overall shape of the toroidal membrane: the side of the toroidal membrane containing FisB had a smaller diameter than the side lacking FisB.

Next, to test if an increase in tension could result in hemi-fission events using pre-stabilized lipids and lipid-replacement of FisB, we increased APL to increase tension to a point where the edges of the membrane did not fill the simulation cube uniformly. In this situation, the solvent filled the remaining gaps at the corners of the simulation cube, which increased lateral membrane tension to at least 14 mN at the toroidal membrane pore region. In these circumstances, the toroidal membrane pore with FisB widened, and after briefly maintaining stability for less than 20 ns, the FisB 20-mer tore apart and led to toroid expansion (Fig. S7Bii, <u>Video 4</u>). These observations suggest that FisB-induced clasping can only tolerate a certain amount of lateral membrane tension. Thus, an increase in lateral membrane tension in the presence of a FisB 20-mer is not sufficient to induce hemi-fission. Importantly, the FisB 20-mer remained stable and did not affect membrane architecture on planar membrane simulations (Fig. S7), suggesting that the effects observed are specific to the toroidal membrane structure. Collectively, these observations suggest that FisB may induce hemi-fission-like events in unstable membranes or act as clasp when membranes are relatively stable.

### Simulation of proteins involved in the final stages division

To establish that the above simulations with FisB are not only artefactual, and demonstrate simulation of other protein assemblies in toroidal membranes using MemTorMD, we ran simulations with SpoIIIE, another bacterial protein thought to exist at a toroidal membrane, but does not play a role in membrane fission (17,39). SpoIIIE, is an FtsK homologue required for DNA translocation into the spore during sporulation and assembles in the polar septum as a single focus during the final stages of division(45,46). SpoIIIE harbours three domains, a membrane domain which includes four membrane-spanning alpha helices, a linker domain and a motor domain (46). The motor domain of SpoIIIE is an ATPase and has been shown to form hexamers *in vitro* which bind DNA and translocate DNA by hydrolysis of ATP (45). Although the oligomerization state of the membrane domain of SpoIIIE and FtsK are still unclear (47), AlphaFold 3 predicts that the SpoIIIE transmembrane domains may form, among other possible homomers, a 12-mer (Fig. S9A) (44). Furthermore, while it is also unclear if this is the configuration of the SpoIIIE transmembrane domain *in vivo,* we nonetheless tested its effects on the toroidal membrane and whether it would also stimulate fission like events in unequilibrated toroids in MemTorMD as observed with FisB. We were able to easily insert the SpoIIIE transmembrane domain 12-mer, without an additional step of lipid replacement, into a 4 nm diameter, 20 nm high toroidal membrane. Interestingly, we noticed that SpoIIIE’s hydrophobic transmembrane regions, due to the symmetrical patterning of the 12-mer, neatly lined the lipid curvature of the lumen of the toroidal membrane pore (Fig. S9). Consistent with the idea that SpoIIIIE is not involved in membrane fission, the toroidal membrane remained intact after 500 ns and the SpoIIIE transmembrane domain 12-mer retained a similar architecture over time (Fig. S9, <u>Video 9</u>). Thus, the introduction of a large membrane-bound oligomeric protein, like the SpoIIIE transmembrane domain 12-mer, does not cause fission-like events and can be simulated in a stable manner.

## DISCUSSION

Computational simulations of complex membrane architectures offer significant potential towards understanding biological problems, especially those where protein function is related to a specific membrane shape. Our work demonstrates that MemTorMD is a versatile and reproducible tool to study toroidal membranes. Not only does our methodology simulate toroidal membranes of different biological dimensions using the triangulated surface to coarse grained framework (TS2CG), it allows the study of lipid behaviour on a coarse grain level in toroidal membranes during nanosecond to microsecond scale simulations. Importantly, MemTorMD can simulate toroidal membranes containing proteins, which has allowed us to gain fundamental insight on a biological event that involves a protein specifically occurring at a toroidal membrane in bacteria. While the work developed here was based on toroidal membranes present within prokaryotic cells, our methodology could also allow for the simulation of eukaryotic toroidal membranes, as it allows inputting lipid composition, concentration and toroidal membrane dimensions.

Previous studies have addressed lipid behaviour and movement on complex membrane architectures, such as membrane buckles, bends and even toroidal membranes (6,8,13–15,37), the approaches used to induce these architectures often relied on artificial features (e.g. force brackets, beads) that constrain lipid movement to a certain shape. While this has enabled insight into the strain and lipid movement present at toroidal membranes induced by these imagined curvatures, and show it is possible to simulate such large lipid assemblies, these studies could not simulate the behaviour of free lipid systems and thus reveal biophysical parameters that enable the stability of toroids in energy minimised states, such as density, size on toroid morphology, or the influence of protein assemblies. Although our determination of membrane architecture intitially relies on a predetermined toroidal shape initially, the lipid stabilization during the simulation leads to an equilibrium within the system which is then not restricted by the initial shape but instead reinforces the shape. Thus, MemTorMD provides a more realistic approach to studying lipid behaviour than existing methods.

Consistent with previous work describing the lipid properties of toroidal membranes of different dimensions and curvature (13), we found that adjusting the APL allowed for stable simulation of toroidal membranes of different sizes and curvature (Fig. 3). Indeed, our simulations of toroidal membranes of different dimensions indicates that APL, and by extension tension, is essential to defining the shape of the toroidal membrane. Notably, we show that APL changes locally within the toroidal membrane, and depends on shape and size, with highly convex curvature causing low density, and highly concave curvature causing high density lipids (Fig. 4). Furthermore, our simulations suggest that we can computationally replicate lipid behaviours that have been observed *in vivo* such as lipid migration (7,24,32,36). We show that cardiolipin typically present in areas of high negative membrane curvature (15), begins to concentrate within the interior leaflet of the pore region of the toroidal membrane which exhibits high negative membrane curvature (Fig. 3C).

One intriguing observation we made with our simulations is that the toroidal membranes, regardless of their dimensions, do not easily fuse on their own. This observation is consistent with the idea that membrane fusion and fission are energetically unfavourable and do not occur spontaneously (16), and that within a biological context membrane fusion and fission is facilitated by proteins (6,8). Based on this, and in the absence of biological data showing known bacterial proteins at toroidal membranes, we used MemTorMD, AlphaFold and planar membrane simulations to study how FisB facilitates the essential fission process that completes the internalization of the developing spore into its adjacent sister cell (24). Consistent with FisB oligomerization observed *in vitro* (7), using MemTorMD we found that FisB can begin oligomerization when randomly inserted around the periphery of the toroidal membrane pore at the start of the simulation (Fig. 5). Interestingly, although the exact configuration of the FisB oligomer at the fission site remains unclear, using AlphaFold2 we found that FisB could form an oligomer composed of 20 FisB molecules (Fig. 5C & D and S6). Co-evolutionary analysis indicates that this FisB 20-mer, or another symmetrical oligomer which has an interface like it such as the 16-mer presented, are likely also physiologically relevant given the position of co-conserved contacting residues, which have been previously implicated in *in-vivo* and *in-vitro* oligomerization, along the oligomeric interface (Fig. S6).

Importantly, and revealing the utility of MemTorMD, in the context of a toroidal membrane we made three important observations that align with what has been observed for FisB *in vitro* and *in vitro* and revealed potential insight into FisB function(7,24). First, in an unstable system with high membrane tension (i.e. when the FisB 20-mer is added during stabilisation at the start of the simulation)(Fig. S8 B, Videos 5-8), hemi-fission like events are observed within the first 100 ns of the simulation, only when FisB is present (Fig. S8). This observation is consistent with a model where FisB-dependent fission requires FisB and increased membrane tension at the narrow neck of the fission pore, which *in vivo* partly occurs by chromosome-induced turgor pressure in the spore compartment (6–8,13,14,24). Second, in a stable system (i.e: when we apply the lipid-replacement method and FisB is added 10 ns into the simulation when membrane tension is low) (Fig. 5E, Fig S7Bi), FisB functions as a clasp and induces asymmetry in the toroidal membrane pore region. This observation is consistent with existing data showing that purified FisB bridges giant unilammelar vesicles and FisB induces small membrane deformities on isolated GUVs that likely represent membrane folds (7). Furthermore, it also suggests that FisB constrains the diameter of the toroidal membrane pore, which may represent an aspect of how FisB induces membrane tension *in vivo*. Third, when high lateral tension is applied to this stable system *in silico*, the FisB 20-mer breaks apart as the pore increases diameter (Fig. S7Bii). This suggests there is a membrane tension threshold after which the FisB oligomers cannot be maintained; it would be interesting to test if this is the case *in vivo.* For example, one could monitor FisB oligomerisation into the fission focus during engulfment, under varying degrees of membrane tension conditions and test if the FisB focus dissociates.

Although we designed MemTorMD to provide a user-friendly interface for studying free toroidal membranes, we recognize some of its limitations. This includes the need to allow stabilization of the toroidal membrane within the first nanoseconds of a simulation, if pore size and stability are a factor in the study of a protein of interest (as we did with FisB), and the semi-automated nature by which one is able to select the protein-insertion point, as well as timing of protein insertion (without an additional lipid-replacement step, MemTorMD requires that the protein be inserted before the start of the simulation). Furthermore, the number of lipids, and complexity of the toroidal membrane architecture, implies that running coarse grain simulations of toroidal membranes is computationally demanding. However, as computational simulations become more widely utilized to study biological problems, and as computational power becomes less limiting, we anticipate that future iterations of MemTorMD-inspired tools will bring accessibility to those looking to investigate protein behaviour on complex membrane architectures. Finally, with the development of new MARTINI 3 beads and models (e.g. the peptidoglycan of bacterial cells) within the Martini-based modelling framework, it should be possible to integrate further complexity into MemTorMD, which will facilitate a greater understanding of protein behaviour within the toroidal membrane occurring during cell division and membrane fission events.

## ACKNOWLEDGEMENTS

We would like to thank the BBSRC for their support of project number BB/X008533/1, awarded to C.D.A.R, which partly funded this work. We would like to thank all members of the Stansfeld and Rodrigues lab for their insights and suggestions, Weria Perezkian for their help in understanding the TS2CG framework software, Sergey Ovchinnikov for the open sharing and instruction of their GREMLIN coevolution GitHub code and Erdem Karatekin, Ane Landajuela and Cecile Morlot for critical reading of the draft manuscript. P.J.S’s laboratory was funded by Wellcome (208361/Z/17/Z), MRC, BBSRC, EPSRC, NIH, JPIAMR and the Howard Dalton Centre. This project made use of time on ARCHER2 granted via the UK High-End Computing Consortium for Biomolecular Simulation, HECBioSim (http://www.hecbiosim.ac.uk), supported by EPSRC (grant no. EP/R029407/1). P.J. S would like to thank the SCRTP at Warwick for use of the computational infrastructure.

## AUTHOR CONTRIBUTIONS

Software writing, experiments, experimental planning, figures and the manuscript was prepared by C. L.B.G, with input, editing and supervision of P.J.S and C.D.A.R. The lipid density tool was created by M.P. Manuscript editing and guidance was provided by P.J.S, M.P and C.D.A.R. Funding acquisition was carried out by C.D.A.R.

## DATA AND MATERIALS AVAILABILITY

All scripts, mdp files and pipelines are available via GitHub or within the Supplementary information. Datafiles containing the calculated APLs, and performed statistics are available in the supplementary .xlx files. Videos of each event are available in

## Supplementary Information

The following files can be found [here](https://figshare.com/articles/dataset/Supplementary_Data_b_Computational_modelling_of_toroidal_membranes_b_b_at_b_b_division_and_fission_sites_b_/28911902/1).

SI1_Figure_3_10us_Domain counting.xlsx

SI2_Figure_4_Cardiolipin chart.xlsx

SI3_Figure_4_Pore_depth_APL.xlsx

SI4_Figure_4_Pore_width_APL.xlsx

SI5_Figure_6_RMSD FisB.xlsx

SI6_Figure_S1_S2_Tension_calculations.xlsx

## Video Files

Video 1. S3A 10usLipiddispersionXY.mp4

Video 2. 5A FisBrandom.mp4

Video 3. S7BI 14nm_500ns_0.6_2STEP_FisB_z.mp4

Video 4. S7BII 14nm_200ns_0.65_2STEP_FisB.mp4

Video 5. S8A FisB20mer_067_XY100ns.mp4

Video 6. S8B FisB20mer_065_XY100ns.mp4

Video 7. S8B FisB20mer_065_Z100ns.mp4

Video 8. S8B FisB20mer_067_XY1us.mp4

Video 9. S9C SpoIIEmem12_XY500nsmov.mp4

---

**Figure S1.**
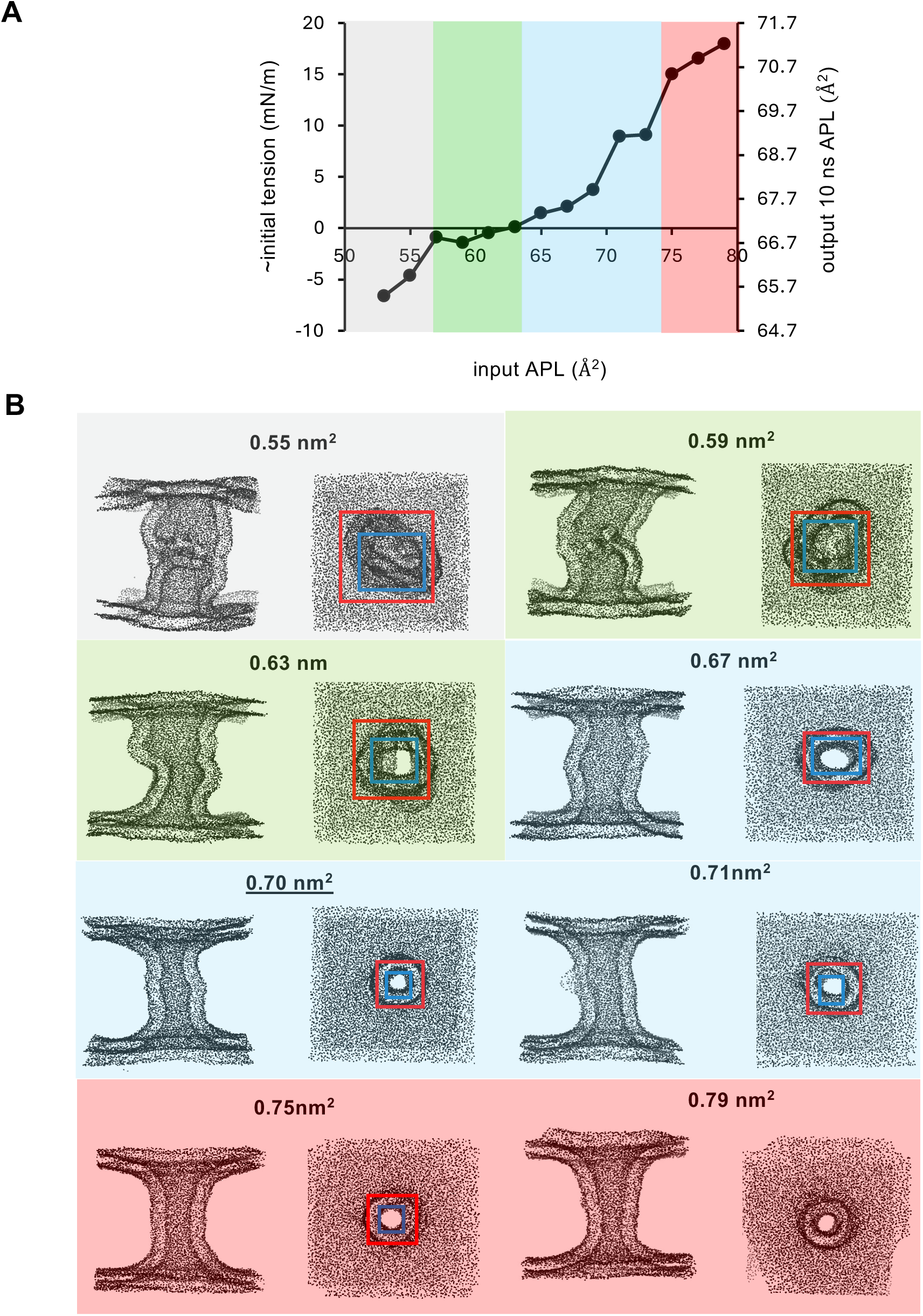
Relationship between input APL and output APL during simulation of a toroidal membrane with 40 nm height and 100% POPG. **(A)** Relationship between tension, input APL and output APL. Colored regions refer to toroidal membranes in different states: unstructured toroidal membranes (grey), zero tension toroidal membranes (green), toroidal membranes under tension (blue) and broken edge toroidal membranes (red). Simulation conditions were 10 ns, 1 bar, 310K, 5e-6 Box, compressibility. **(B)** Examples of toroidal membranes in different states as described in (A); Phosphates displayed as spheres. Z 40 nm, XY 4 nm; blue and red box represent inner and outer pore dimension measurements, respectively. Input APL is shown above each example.

**Figure S2.**
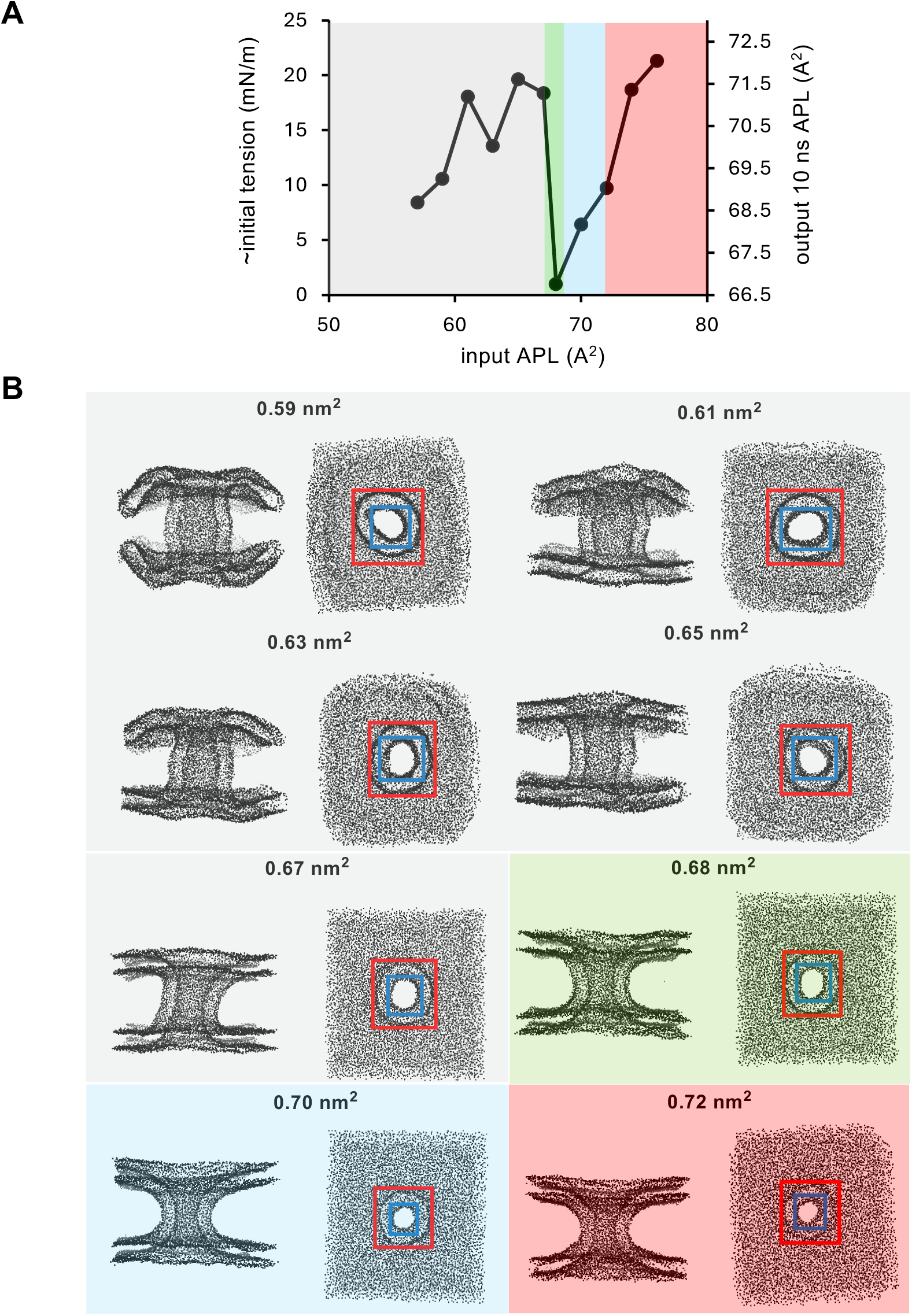
Relationship between input APL and output APL during simulation of a toroidal membrane with 20 nm height and 100% POPG. **(A)** Relationship between tension, input APL and output APL. Colored regions refer to toroidal membranes in different states: unstructured toroidal membranes (grey), zero tension toroidal membranes (green), toroidal membranes under tension (blue), and broken edge toroidal membranes (red). Simulation conditions were 10 ns, 1 bar, 310K, 5e-6 Box, compressibility. (**B)** Examples of toroidal membranes in different states as described in (A); Phosphates displayed as spheres. Z 20 nm, XY 4 nm; blue and red box represent inner and outer pore dimension measurements, respectively. Input APL is shown above each example.

**Figure S3.**
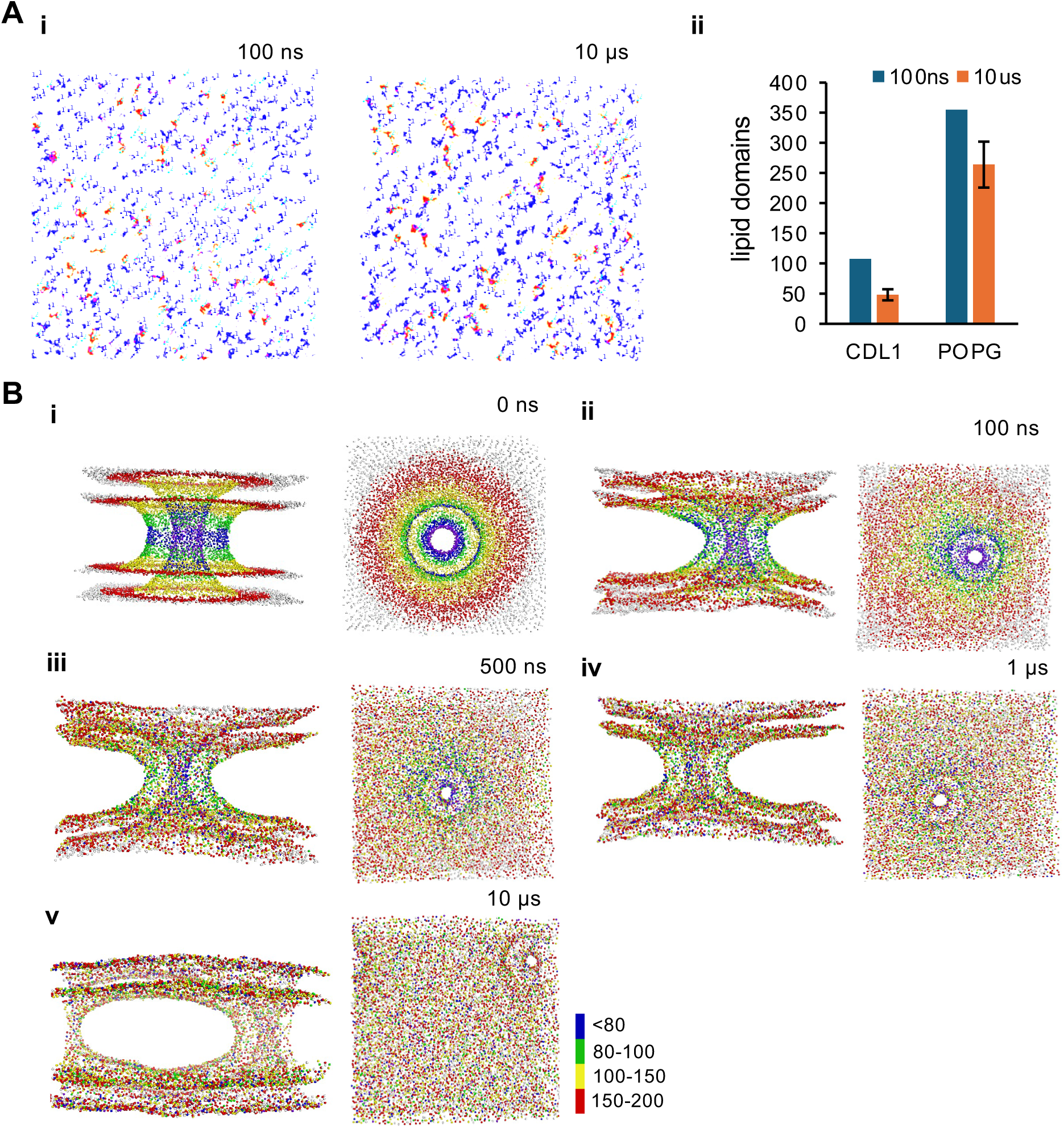
Lipid mixing and lipid domains during simulation of toroidal membranes in MemTorMD. **(A)** Lipid domain visualization and frequency determination; simulation conditions were POPE 78%, POPG 20%, CDL1 2%, 310k, 1 bar; POPG (blue), CDL1 (orange). **(i)** Fixed starting value 100 ns 355 POPG clusters, 108 CDL1 clusters after 10 µs. POPE hidden **(ii)** Manual lipid domain frequency determination after 10 µs, compared to fixed position at the start of simulation. (**B)** Lipid mixing over time. Lipid distance from center of the toroidal membrane is colour-coded to illustrate lipid mixing; red 200 Å, yellow 150 Å, green 100Å and blue 80 Å.

**Figure S4.**
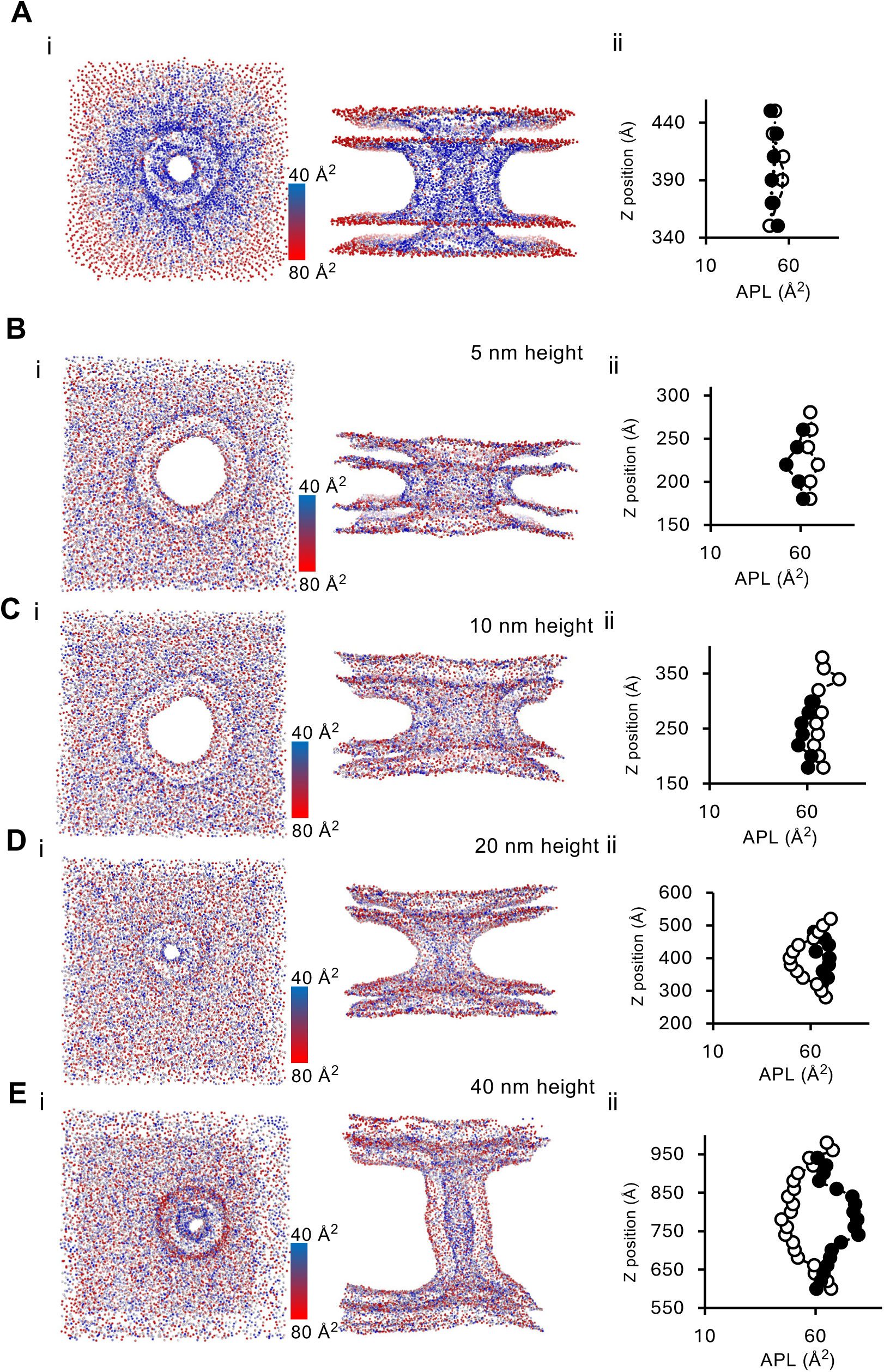
Relationship between local APL within the cylindrical region (Z-plane) of the toroidal membrane, as a consequence of toroidal membrane height. **(A-E)** APL values at different Z position averages along the toroidal membrane: during coarse-grained simulation of toroidal membranes with different height. (**A)** APL distribution at simulation start (0 ns) in a 20 nm high toroidal membrane. (**B-E**) APL distribution after simulation (0.5 µs) in toroidal membranes of different height. Simulation conditions were POPE 78%, POPG 20%, CDL1 2%, at 310K, 1 bar. (i) visualization of APL using a colour-coded phosphate representations (see scale - scale is represented by tones of red and blue where red is 80 Å^2^ APL and blue is 40 Å^2^ APL); (ii) Average APL (Å^2^) at different Z-positions along the toroidal membrane cylinder.

**Figure S5.**
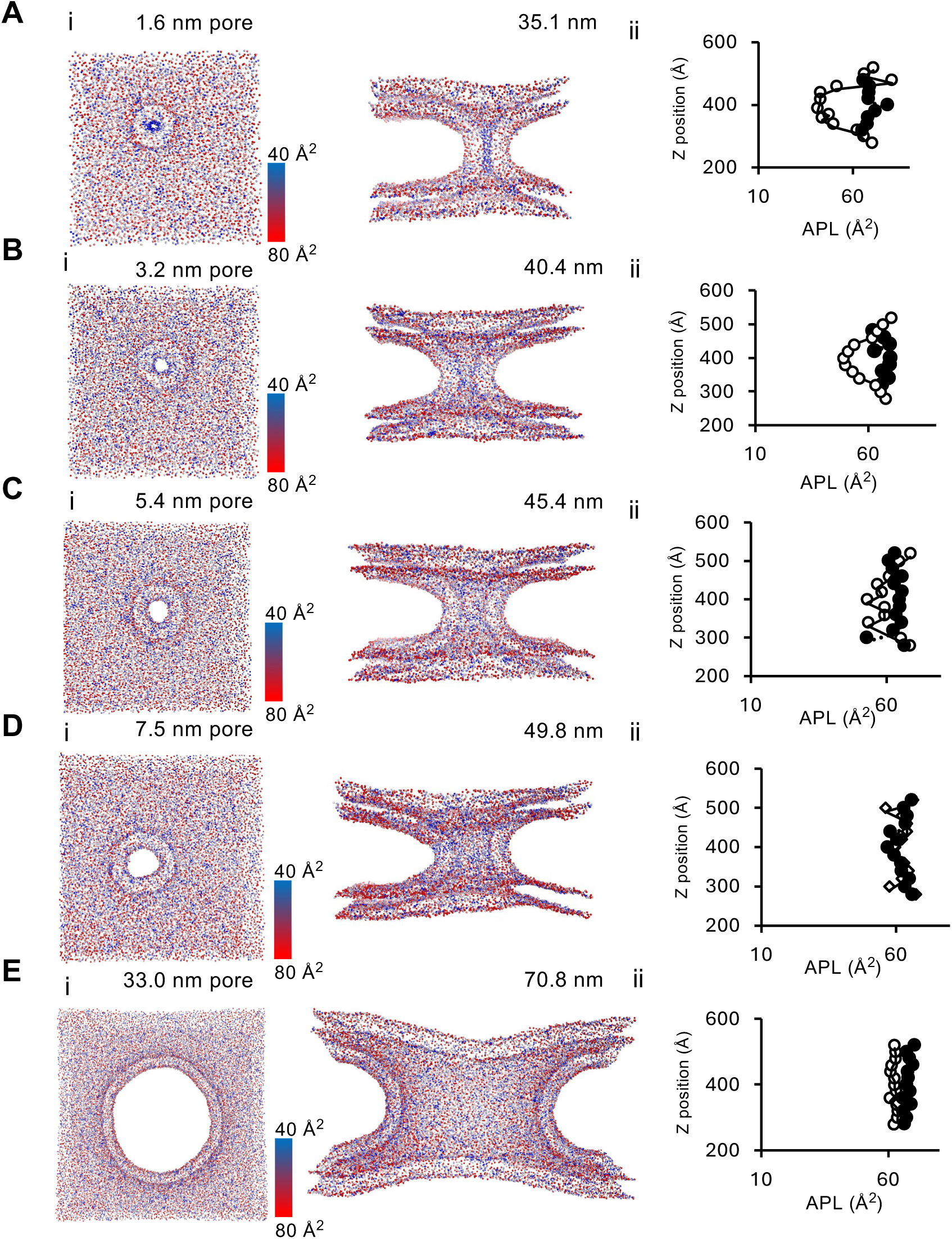
Relationship between local APL within the cylindrical region (Z-plane) of the toroidal membrane, as a consequence of toroidal membrane pore diameter. **(A-E)** APL values at different Z position averages along the toroidal membrane: during coarse-grained simulation of toroidal membranes with different height. (**A-E**) APL distribution after simulation (0.5 µs) in toroidal membranes of different pore diameter. Simulation conditions were POPE 78%, POPG 20%, CDL1 2%, at 310K, 1 bar. (i) visualization of APL using a colour-coded phosphate representations (see scale - scale is represented by tones of red and blue where red is 80 Å^2^ APL and blue is 40 Å^2^ APL); (ii) Average APL (Å^2^) at different Z-positions along the toroidal membrane cylinder.

**Figure S6.**
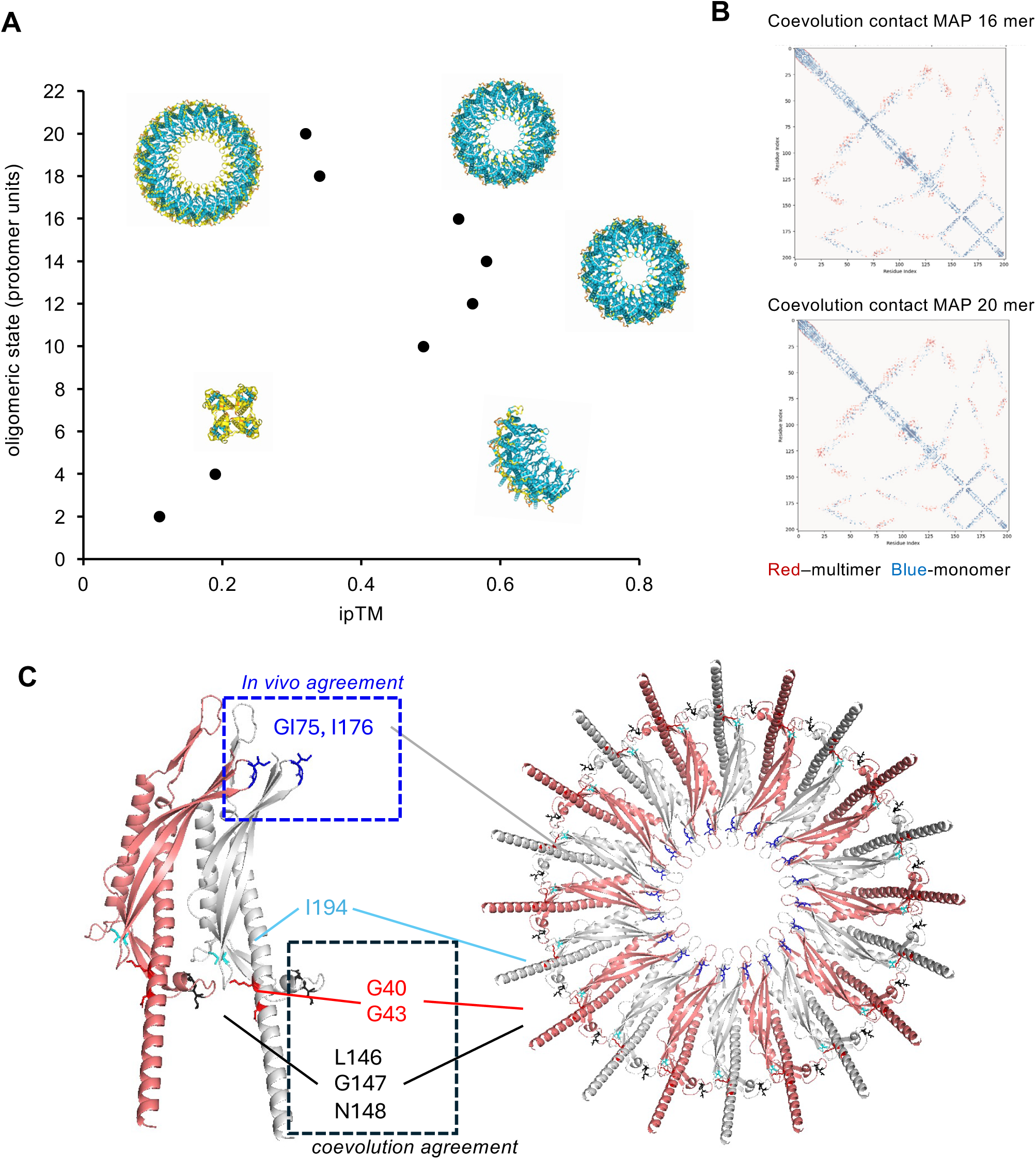
FisB multimerization based on AlpaFold3 and coevolutionary analysis of FisB contacts within multimers. **(A)** FisB multimer comparisons by ipTM score predicted by AlphaFold3. (**B)** Coevolution matrix (GREMLIN) comparing “FisB multimer” (red) to “FisB monomer” (blue). Comparisons for the FisB 20-mer and 16-mer are shown. (**C)** FisB 20-mer interface showing residues implicated in FisB oligomerization *in vivo,* G175,176 (blue box). I194 (cyan) is also implicated in FisB oligomerization *in vivo* but is not predicted by co-evolution to establish contacts between protomers. Residues L146, G147 and N148 (black) are predicted by co-evolution to interact with G40 and G43 (red) and lie the FisB-20 mer protomer interface.

**Figure S7.**
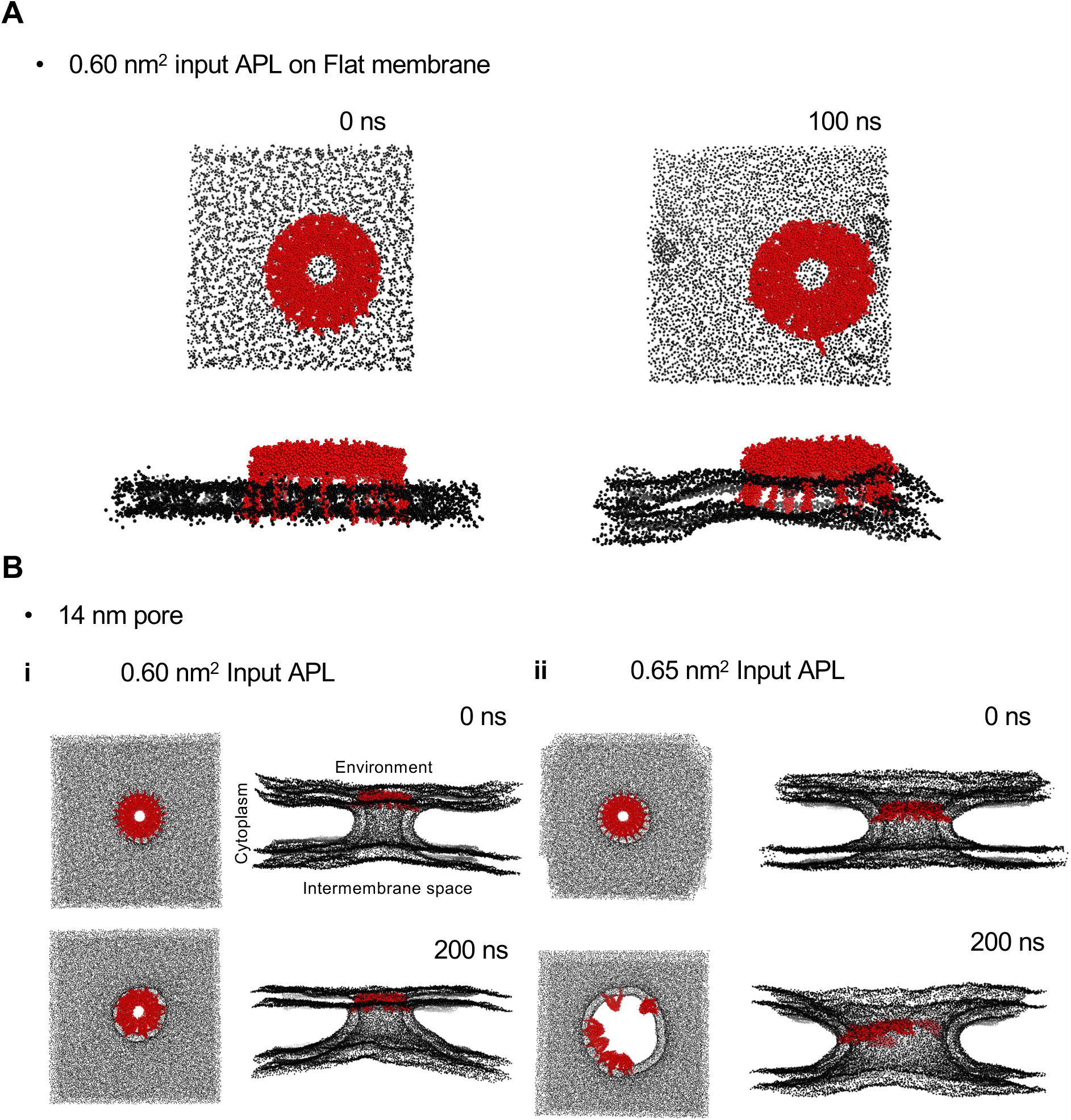
Simulation of FisB on a flat membrane and comparison of FisB 20-mer dynamics at a pre-stabilized toroidal membrane. **(A)** Hypothetical FisB 20mer dynamics after insertion by TS2CG methodology on flat membrane for 100 ns, 1bar, 310K, POPE 78%, POPG 20%, CDL1 2% (**B)** FisB 20-mer dynamics at 200 ns after insertion by lipid replacement on toroidal membranes pre-equilibrated for 10 ns: (**i)** Low tension (0.60 nm^2^ Input APL), showing a clasp-like event and asymmetric toroid (**ii)** High tension (box edge breakage; 0.65 nm^2^ input APL) causes FisB 20-mer dispersal. Input toroidal membrane pore diameter was 14 nm input with POPE 78%, POPG 20%, CDL1 2%.

**Figure S8.**
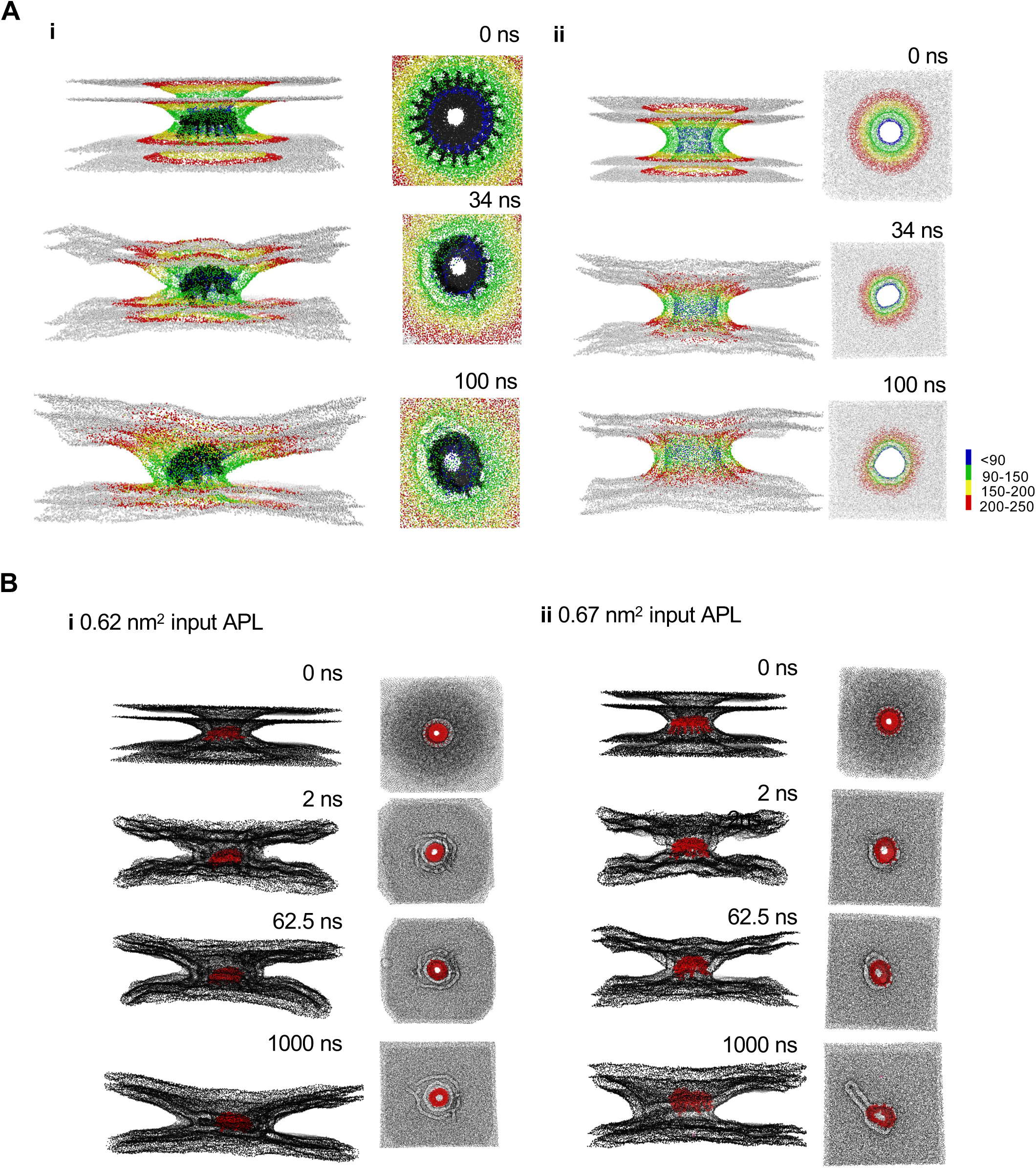
Lipid movement analysis during hemi-fission-like events caused by FisB 20-mer. **(A)** Lipid distribution overtime during 0.67 nm^2^ input APL hemi-fission event. “Lipid distance from center of the toroidal membrane” is color-coded to illustrate lipid mixing; red – 200 Å, yellow 150 Å, green - 100Å and blue - 80 Å. Little lipid mixing occurs **(i)** with FisB 20-mer and (**ii)** no FisB 20-mer control. **(B)** Simulation of FisB 20-mer without pre-equilibration can cause hemi-fission like events in a toroidal membrane with 18 nm pore diameter; Lipids concentration during simulation were: POPE 78%, POPG 20%, CDL1 2%.

**Figure S9.**
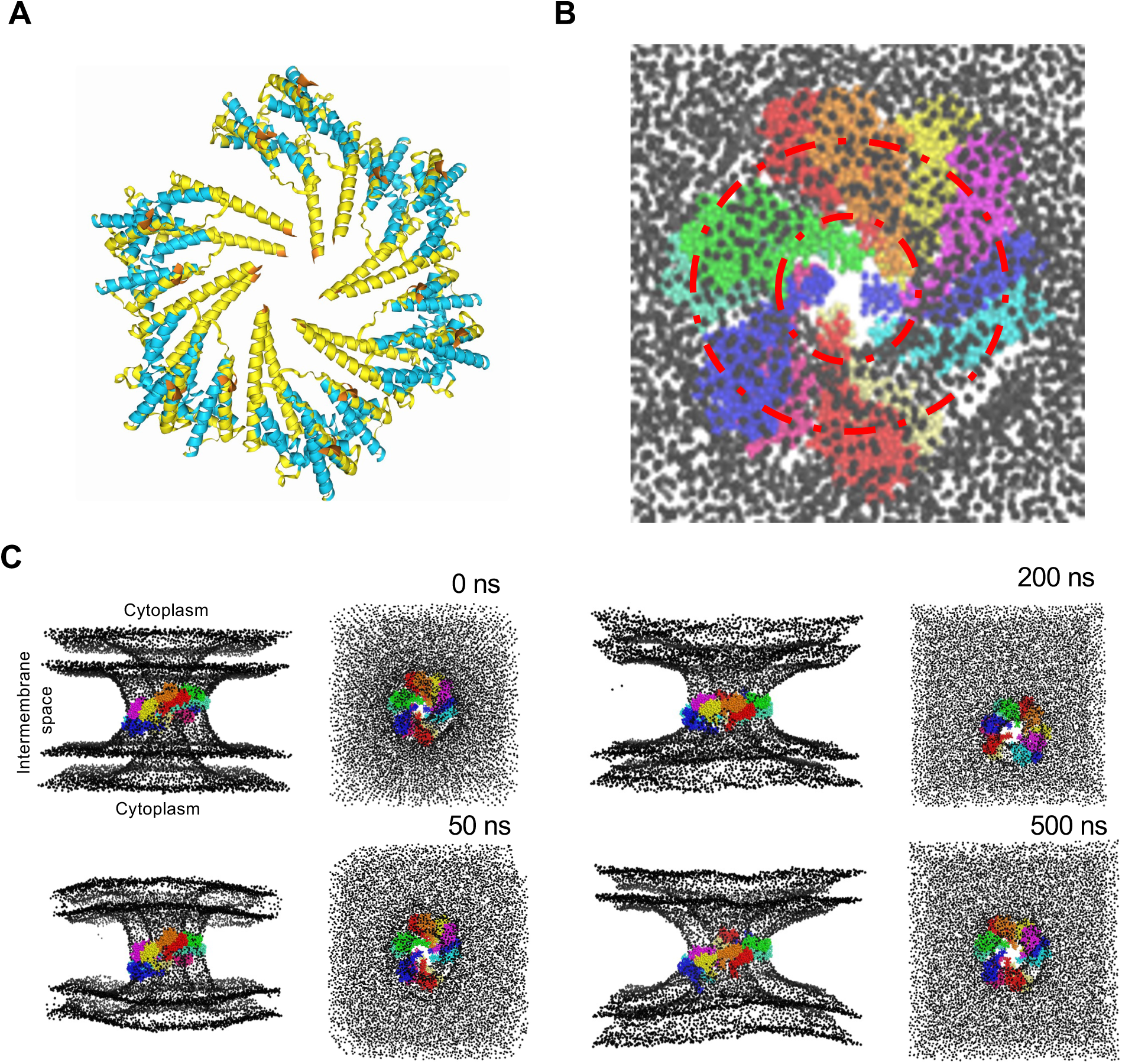
Stable insertion of the SpoIIIE transmembrane domain 12-mer into a toroidal membrane pore. **(A)** AlphaFold 3 predicted structure of SpoIIIE transmembrane 12-mer; LddT indicated by colour. **(B)** Predicted SpoIIIE transmembrane 12-mer insertion into a 3.2 nm toroidal membrane pore by MemTorMD script. Red dashed line indicates toroidal membrane pore boundaries. **(C)** Simulation of SpoIIIE transmembrane 12-mer; toroidal membrane was set to 0.65 nm^2^ input APL and 3.2 nm pore width and simulation for over 500 ns. Lipid composition was POPE 78%, POPG 20%, CDL1 2%.

